# PySCNet: A tool for reconstructing and analyzing gene regulatory network from single-cell RNA-Seq data

**DOI:** 10.1101/2020.12.18.423482

**Authors:** Ming Wu, Tim Kacprowski, Dietmar Zehn

**Affiliations:** Division of Animal Physiology and Immunology, School of Life Sciences Weihenstephan, Technical University of Munich, Freising, 85354, Germany; Research Group on Computational Systems Medicine, Chair of Experimental Bioinformatics, TUM School of Life Sciences, Technical University of Munich, Freising, Germany; Division of Data Science in Biomedicine, Peter L. Reichertz Institute for Medical Informatics of TU Braunschweig and Hannover Medical School, Brunswick, Germany

## Abstract

**Summary:** The Advanced capacities of high throughput single cell technologies have facilitated a great understanding of complex biological systems, ranging from cell heterogeneity to molecular expression kinetics. Several pipelines have been introduced to standardize the scRNA-seq analysis workflow. These include cell population identification, cell marker detection and cell trajectory reconstruction. Yet, establishing a systematized pipeline to capture regulatory relationships among transcription factors (TFs) and genes at the cellular level still remains challenging. Here we present PySCNet, a python toolkit that enables reconstructing and analyzing gene regulatory networks (GRNs) from single cell transcriptomic data. PySCNet integrates competitive gene regulatory construction methodologies for cell specific or trajectory specific GRNs and allows for gene co-expression module detection and gene importance evaluation. Moreover, PySCNet offers a user-friendly dashboard website, where GRNs can be customized in an intuitive way.

**Availability:** Source code and documentation are available: https://github.com/MingBit/PySCNet

## 1 Introduction

An increasing number of software packages and pipelines have been developed for interpreting high content gene expression data generated through the utilization of high throughput single-cell sequencing technologies (Satija *et al.*, 2015; Kiselev *et al.*, 2017; Qiu *et al.*, 2017; Wolf *et al.*, 2018). These tools predominantly aim to investigate cellular subsets distributions and spatial diversities via clustering and cell trajectory construction. Moreover, there are multiple tools that allow us to identify and compare cluster specific differential gene expression patterns. However, scRNA-seq datasets harbour unpreceded further information about gene co-expression networks and molecular dependencies that are, given the lack of powerful tools, rarely explored. So far, several methods have been proposed to capture gene-gene or gene-TFs interactions in the last decade (Huynh-Thu *et al.*, 2010; Chan *et al.*, 2017; Matsumoto *et al.*, 2017; Woodhouse *et al.*, 2018 and Moerman *et al.*, 2019), but the outcomes yielded from different methods are largely irreconcilable (Chen and Mar, 2018). Also, it is challenging to evaluate the results in the absence of widely confirmed ground truth. We therefore developed PySCNet, a python package which provides diverse network inference methods to offer more flexibility for gene regulatory networks (GRNs) reconstruction. Integrating graph-based approaches, our package extends network analysis to facilitate a better understanding of molecular networks.

## 2 Methods

To investigate the molecular characters from single-cell RNA-seq data beyond the current toolkits, PySCNet takes the results from other toolkits and builds cell specific GRNs. Meanwhile, it provides graph based techniques to summarize the strength of regulatory interactions, which can be uploaded onto an interactive web-based dashboard.

**Fig. 1.**
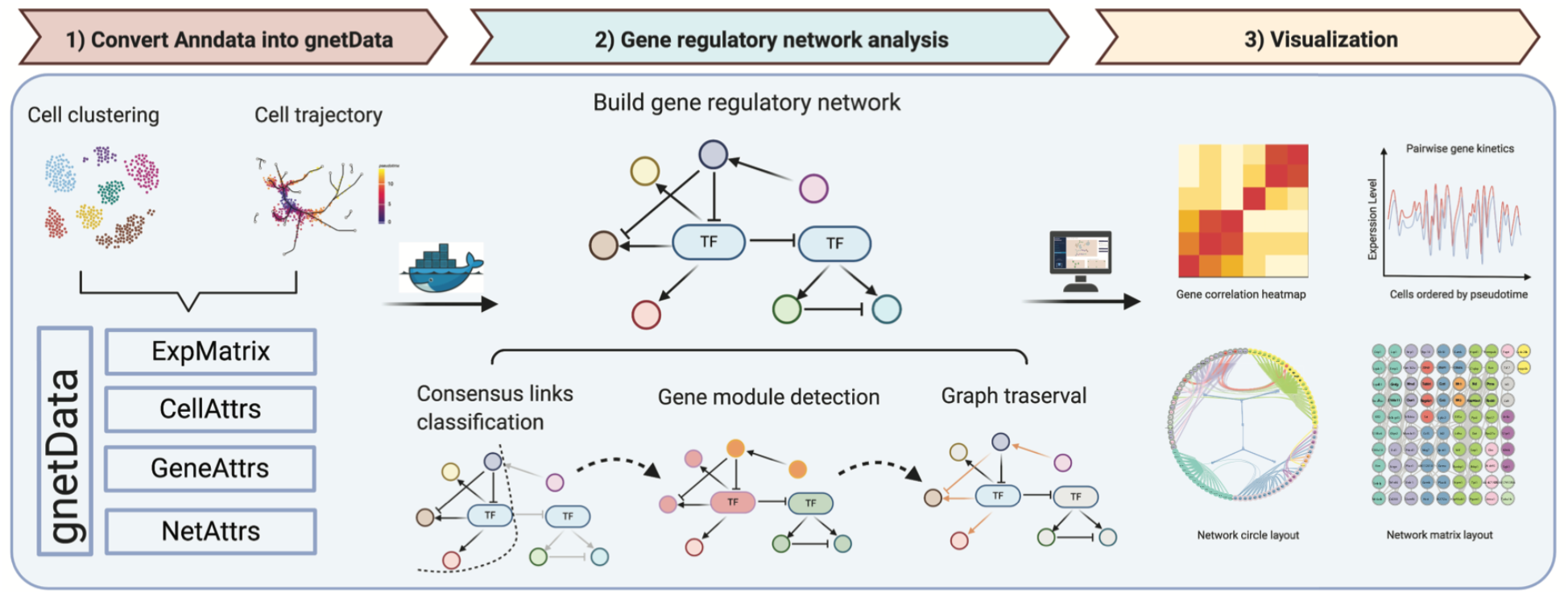
Overview of PySCNet workflow. 1) Single cell mRNA expression matrices with cell / gene annotations stored in AnnData are converted into gnetData. 2) Integrating gene inference methods with Docker, GRNs are reconstructed followed by network analysis including consensus regulation prediction, gene module detection and gene path traversal. 3) The correlation heatmap and gene kinetic plot are illustrated for subnetwork and the global network can be viewed on PySCNet dashboard in different layouts.

### 2.1 PySCNet provides an interface to other toolkits built on AnnData

PySCNet introduces a python class for gene network annotation – gnetData. The results obtained from widely used toolkits such as scanpy (Wolf *et al.*, 2018) and stream (Chen *et al.*, 2019), can be directly imported into PySCNet.

### 2.2 PySCNet integrates competitive GRN inference methods

To infer gene regulatory interactions, we selected GENIE3 (Huynh-Thu *et al.*, 2010), PIDC (Chan *et al.*, 2017) and GRNBOOST2 (Moerman *et al.*, 2019), which are concluded as competitive tools in terms of reconstructing GRNs from curated models and experimental datasets (Pratapa *et al.*, 2020). Furthermore, we added sliding window correlation (SWC) and phase synchrony measurement for gene co-expression analysis. PySCNet integrates Docker to build GRNs applying the aforementioned competitive regulatory inference methods.

Networks yielded from different methods can be merged, or classified via binary classification algorithms. We used Louvain clustering to detect gene co-expression modules. To determine master TFs dominantly regulating other genes, node centrality is applied to estimate the importance of gene / TF in the network. In order to discover hidden regulating links of a target gene node, graph traversal techniques are utilized to predict indirect regulations.

### 2.3 PySCNet offers a Dashboard to interactively customize gene network

As arbitrary thresholds or parameters are involved in constructing networks, it is necessary to tune the parameters to get optimal networks. To keep this practical, we designed a simple dashboard with parameter settings, which allows for parameters adjustment.

## 3 Case study

To demonstrate the module usage of PySCNet, we extended cell clustering analysis on peripheral blood mononuclear cells (PBMCs) guided by scanpy (https://scanpy-tutorials.readthedocs.io/en/latest/pbmc3k.html). We applied GENIE3, PIDC, GRNBOOST2 and Phase synchrony to build cell specific gene networks on B, CD4 T, CD8 T and CD14 Monocytes (Supplementary Fig.1). Node centrality was measured and ranked to evaluate the importance of the gene in the network. This analysis demonstrates that the highest ratio of rank score of CD14, IL7R, CD8A were found in CD14 monocytes, CD4 T cells and CD8 T cells, which is accordant with expression profiles. Our results further confirm the B-cell specific TF (SPIB), T-cell specific TFs (LYAR, HOPX) and shared TFs (IRF8, SPI1, CEBPD) (Supplementary Fig.2).

We further performed cell trajectory analysis on mouse hematopoietic stem cells (HSCs) (Nestorowa et al., 2016) guided by stream. To track how genes co-expressed along the individual cell lineage, 535 cells assigned to the megakaryocyte-erythrocyte progenitors (MEP) cell line and 236 highly expressed genes were selected for GRN construction applying 6 methods (Supplementary Fig.3). We found that pairwise gene interaction ranked by phase synchrony gives an optimal outcome in terms of early precision evaluation with top 420 edges (Supplementary Fig.4). In furtherance of network illustration, a pickle object derived from PySCNet was uploaded onto the dashboard, which gives rise to more flexible options of network illustration (Supplementary Fig.7).

## 4 Conclusion

PySCNet accommodates an extensive variety of GRN construction algorithms, graph based techniques and a user-friendly interface to investigate regulatory relationships among genes and transcription factors from single-cell RNA-seq data. Instead of following the mainstream analysis on cell behaviour, PySCNet opens a window to systematically inspect gene-gene / TF communication at single cell resolved level.

## Acknowledgements

We gratefully acknowledge useful discussion with Alex Hildebrandt.

