## Supplemental Data for "PySCNet: A tool for reconstructing and analyzing gene regulatory network from single-cell RNA-Seq data"

### Supplementary Note

To exemplify the usage of PySCNet, we provide two case studies including extending cell clustering analysis of Peripheral Blood Mononuclear Cells (PBMC) guided by scanpy [1] and cell trajectory analysis of mouse Hematopoietic Stem Cells (HSC) guided by stream [2].

##### Case study: B-cell / T-cell specific transcription factors are identified in PBMC data

The PBMC data is freey accessiable from 10X genomics (`https://support.10xgenomics.com/single-cell-gene-expression/datasets/1.1.0/pbmc3k`). Cells were grouped into 8 sub-types and were labelled by the known markers (Figure 1.A). In order to build cell specific gene networks, we took cell populations with abundance greater than 200 and the union set ($\sim 190$) of top 50 highly expressed genes derived from selected populations (Figure 1.C). GENIE3 [3], PIDC [4], GRNBOOST2 [5] and Phase Synchrony were applied for network inference. Among those methods, the overlapping set ($\sim 8900$) of top 15000 interactions achieved by individual methods were selected for graph construction (Figure 1.B).

With regard to identifying the master transcription factors dominantly regulating other genes, node centralities including degree, closeness, betweenness and pagerank were calculated and ranked. The average of rank score was used to evaluate the importance of the node in the network. With respect to the percentage of rank score in each cell types, we observed that the highest ratio of rank score of CD14, IL7R, CD8A were found in CD14 monocytes, CD4 T cells and CD8 T cells (Figure 2.A, 2.B), which is consistent with expression profiles. This observation is of particular interest in terms of predicting cell specific transcription factors. Indeed, our results highlight the B-cell specific transcription factor (SPIB), T-cell transcription factors (LYAR, HOPX) and shared transcription factors (IRF8, SPI1), which have been previously reported [6, 7, 8, 9, 10, 11, 12].

The code is available at `https://github.com/MingBit/PySCNet/blob/master/tutorial/pyscnet_scanpy.ipynb`

**Case Study: Phase Synchrony outperforms other GRN methods in terms of early prediction in mouse HSC data**

Mouse HSC data obtained from [13] were re-analyzed by stream. Cells were divided into six sub-groups and were formed into two distinct cell lines. Hematopoietic stem cells (HSCs) expanded towards: 1) megakaryocyte-erythrocyte progenitors (MEPs); 2) the mixture of lymphoid-primed multipotent progenitors (LPMPs) and granulocyte-monocyte progenitors (GMPs).

In order to examine the relationships among the leaf markers along the MEP branch, we selected 535 cells assigned to the MEP cell line and 236 positive leaf markers to build gene network. Protein-protein interactions downloaded from stringPPI [14] were considered as reference for performance evaluation. 420 edges were filtered by high confidence scores greater than 0.7 and enrichment p-value smaller than 1.0e-16. As the reference network was connected by 117 features, other 119 isolated nodes without connection were removed for GRN construction. Top 420 edges were selected from each method for early precision evaluation. As shown in figure 4, while GENIE3, GRNBOOST2, PIDC and Window sliding based correlation give comparable results, phase synchrony based method outperforms other algorithms.

Moreover, we found that interactions excluded from the top list, were identified by greedy walk. For instance, connections including Mki67-Rrm2, Mki67-Mcm5, Mki67-Mcm7 failed to be predicted but surprisingly found in the 5 steps of greedy walk.

The code is available at `https://github.com/MingBit/PySCNet/blob/master/tutorial/pyscnet_stream.ipynb`

### Supplementary Data

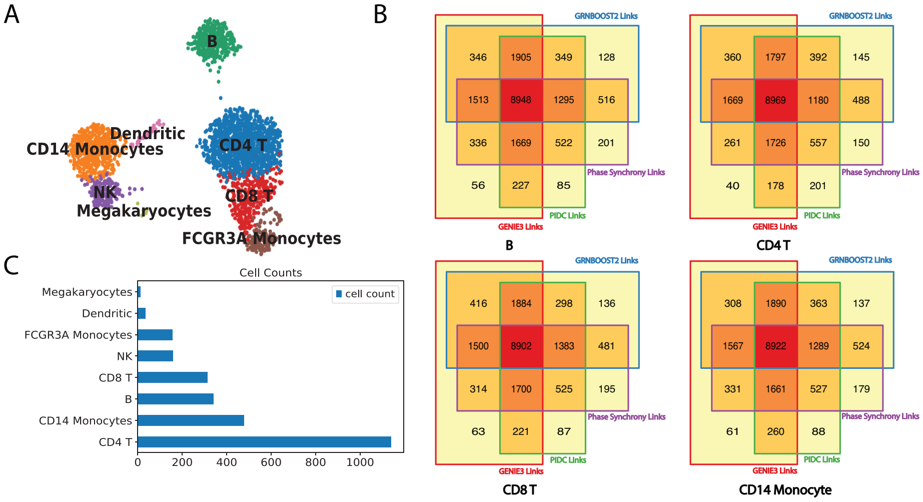


Figure 1: **PBMC cell clustering and cell specific gene networks.** A) 8 cell types distincted by leiden clustering [15] and labelled by known markers. B) The bar plot showing cell abundance in the groups. C) Venn diagrams illustrating consensus regulation links of top 15000 interactions derived from GENIE3, PIDC, GRNBOOST2 and Phase Synchrony.

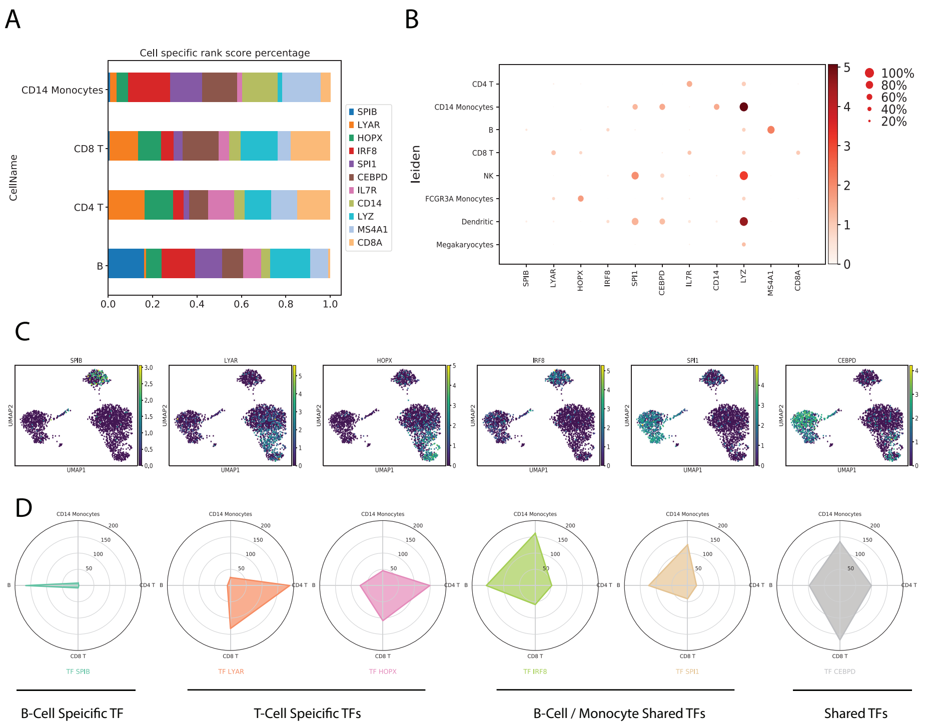


Figure 2: **Cell specific and shared TFs were identified by ranking node centrality.** A) The percentage of rank score indicating cell markers have the highest ratio in the associated cell populations. B) The dotplot denoting gene expression in cell groups. Circle size is encoded by the fraction of cells in the group and color is encoded by the mean expression in the group. C) Feature plots and radar plots illustrating the cell specific and shared transcription factors.

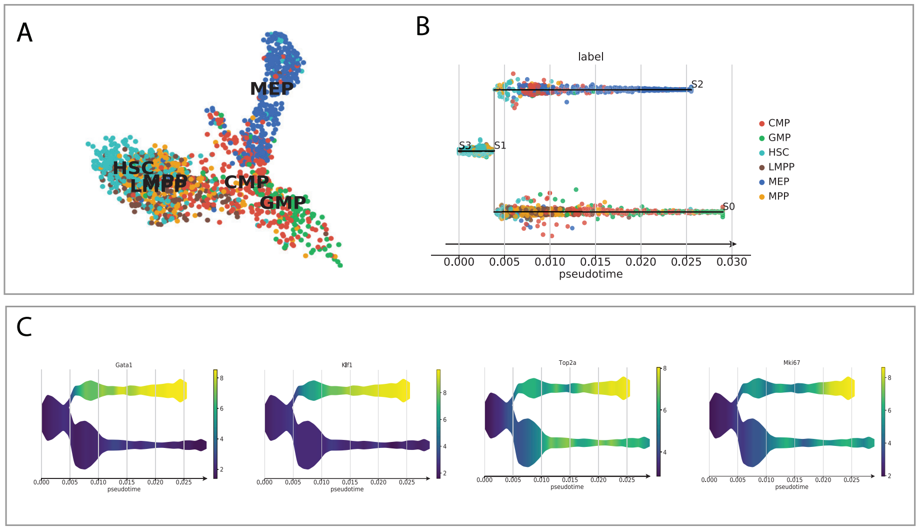


Figure 3: **Cell clustreing and development trajectory of Mouse hematopoietic stem cell**. A) 6 sub-populations labelled and illustrated in Uniform Manifold Approximation and Projection (UMAP) by scanpy [1]. B) Hematopoietic stem cells (HSCs) differentiate into two cell lines. C) 4 representative Leaf markers (Gata1, Klf1, Top2a and Mki67) are highly expressed in MEP branches.

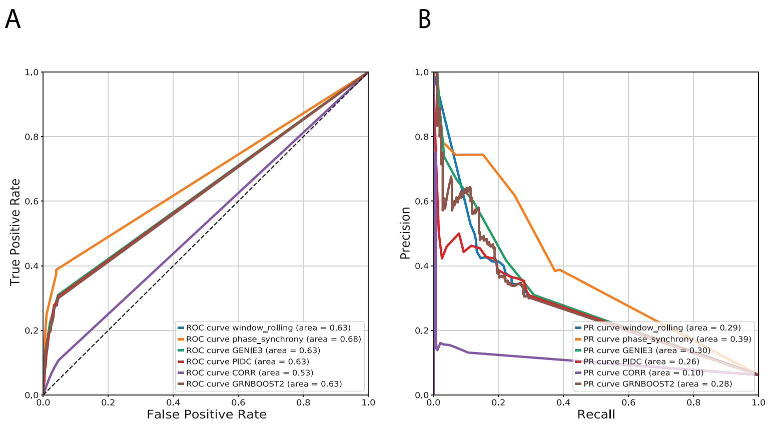


Figure 4: **Algorithms performance.** Gene regulatory networks (GRNs) were generated from 6 methods. Reference links were downloaded from string PPI and top 420 edges were selected for performance evaluation A) Receiver Operating Characteristic (ROC) curve for top 420 edges. B) Precision Recall (PR) curve for top 420 edges.

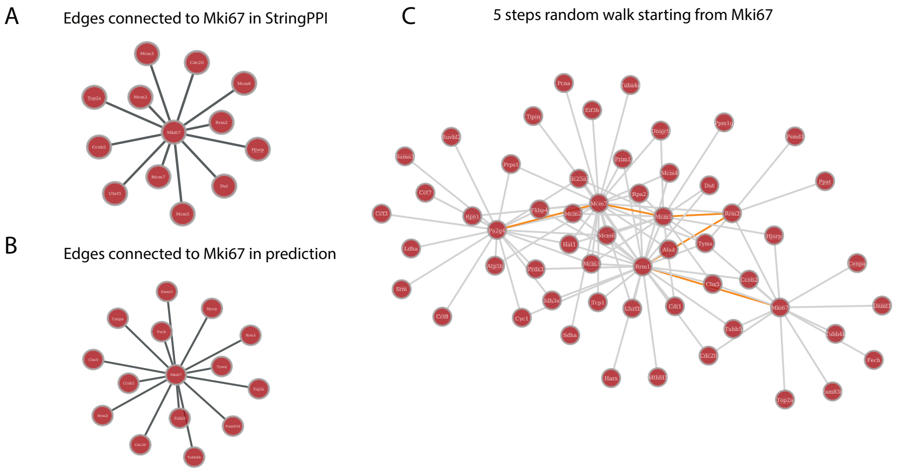


Figure 5: **Explore gene network via greedy walk.** A) Reference edges connected to Mki67 in stringPPI filtered by high confidence score $> 0.7$ and enrichment p-value $< 1.0e\text{-}16$. B) Top edges connected to Mki67 predicted by phase synchrony method. C) Sub-network shows the 5 steps greedy walk guided by node degree starting from Mki67: (Mki67 - Rrm1 - Rrm2 - Mcm5 - Mcm7 - Pa2g4).

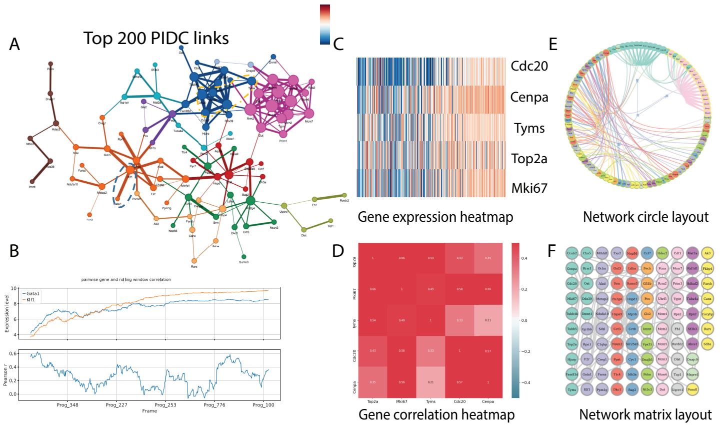


Figure 6: **Gene network visualization.** A) Top 200 links obtained from PIDC. Nodes were colored by gene modules and edges were weighted by interaction strength. B) Pairwise gene kinetics (Gata1-Klf1) along the cell trajectory. C, D) 5 nodes sub-network (Cdc20, Cenpa, Tyms, Top2a and Mki67) was illustrated as expression heatmap and correlation heatmap. E,F) The global view of the network was shown in circle and matrix layout.

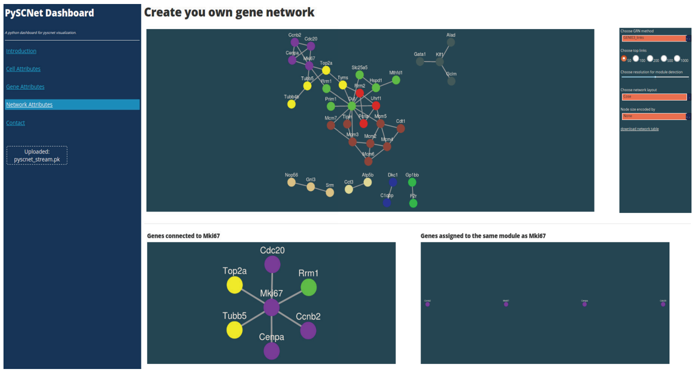


Figure 7: **PySCNet-Dashboard.** A pickle object derived from PySCNet was uploaded onto PySCNet-Dashboard, which allows for parameters adjustment to interactively optimize gene networks.
